# FinDonor 10 000 study: A cohort to identify iron depletion and factors affecting it in Finnish blood donors

**DOI:** 10.1101/507665

**Authors:** Lobier Muriel, Niittymäki Pia, Nikiforow Nina, Palokangas Elina, Larjo Antti, Mattila Pirkko, Castrén Johanna, Partanen Jukka, Arvas Mikko

## Abstract

**Background and Objectives:** There is increasing evidence that frequent blood donation depletes the iron stores of some blood donors. The FinDonor 10 000 study was set up to study iron status and factors affecting iron stores in Finnish blood donors. In Finland, iron supplementation for at-risk groups has been in place since the 1980’s.

**Material and Methods:** 2584 blood donors (N= 8003 samples) were recruited into the study alongside standard donation at three donation sites in the capital region of Finland between 5/2015 and 12/2017. All participants were asked to fill out a questionnaire about their health and lifestyle. Blood samples were collected from the sample pouch of whole blood collection set, kept in cool temperature and processed centrally. Whole blood count, CRP, ferritin and sTFR were measured from the samples and DNA was isolated for GWAS studies.

**Results:** Participant demographics, albeit in general similar to the general blood donor population in Finland, indicated some bias toward older and more frequent donors. Participation in the study increased median donation frequency of the donors. Analysis of the effect of time lag from the sampling to the analysis and the time of day when sample was drawn revealed small but significant time-dependent changes.

**Conclusion:** The FinDonor cohort now provides us with tools to identify potential donor groups at increased risk of iron deficiency and factors explaining this risk. The increase in donation frequency during the study suggests that scientific projects can be used to increase the commitment of blood donors.

## Introduction

Approximately 200 - 250 mg of iron is drawn in a standard blood donation; this amount accounts for 25% of average tissue iron stores in men and up to 75% in women. There is compelling evidence that a portion of blood donors may become iron depleted or deficient. Deferral rates of presenting donors, due to too low level of hemoglobin, vary a lot depending on populations and policies from below 10 % to up to 20 %. In addition, for example, in the U.S. 35% of frequent blood donors were found to be iron deficient [1–6]. Adverse health effects with various symptoms such as fatigue, pica, restless legs syndrome, and cognitive problems have been linked to iron deficiency and anemia [7], although the effect of iron deficiency without anemia in otherwise healthy individuals is still unclear [8]. The iron removed by blood donation should be replaced by dietary iron. Another tool to ensure correction of iron stores is the minimum time interval between blood donations, which must be balanced between donor health issues and blood demand. There is evidence [2,4,9] that the current intervals may not be sufficient for iron or Hb recovery at least for some donors. Identification of donors who are at highest risk for developing iron depletion or anemia as a result of frequent blood donation is warranted as it could give us fact-based tools to steer donor recruitment.

Implementation of iron supplementation to an entire blood donor population to manage iron deficiency is rare, although several countries provide some iron supplementation [10]. The Finnish Red Cross Blood Service (FRCBS) has implemented a national risk-group based iron supplementation policy since the 1980’s. Iron tablets are provided at donation time to all female donors under the age of 50 and to all other donors donating every 4^th^ month or more frequently.

To complement the above mentioned studies on iron depletion and cohorts collected elsewhere [11–14], we set up a cohort study focusing primarily on the measurement of iron stores, factors affecting them, and, in addition, on the effect of systematic iron supplementation. The study had 3 specific goals:

1. To assess the feasibility of carrying out research embedded in the regular blood donation process of FRCBS.
2. To assess the current status of iron biomarkers in the Finnish blood donor population and their relationship to self-reported health and genetic background.
3. To provide preliminary data on the evolution of iron biomarkers for estimating sample sizes of future studies.

In consecutive publications, we will analyze distributions of iron biomarkers and estimate their determinants with multivariate regression (goal 2) [15]. We will subsequently carry out genome wide association analysis for iron biomarkers. We will use results from these studies to carry out simulations for estimating sample-sizes for future studies (goal 3).

Based on our sociological study, Finnish blood donors are willing to donate also for research use [16]. In general, blood donors have been regarded as an important resource for biomedical studies [17]. We here describe the FinDonor 10 000 cohort, its collection process, possible sources of bias and give evidence that there are factors in the measurements that may affect the results if not carefully taken into account.

## Materials and methods

The Finnish Red Cross Blood Service (FRCBS) is a national blood establishment responsible for the collection, processing and distribution of blood products in Finland. An Ethical review and approval for the study was obtained from the Ethical Board of Helsinki University Hospital, Helsinki, Finland. Informed consent was obtained from participants and they were allowed to withdraw from the study at any time.

### Sample collection

The FinDonor 10 000 study ran from May 18th 2015 to December 8th 2017. All donors in the participating three fixed donation sites located in the Helsinki metropolitan area (Kivihaka, Sanomatalo, Espoo) were informed of the possibility to join the study with leaflets, posters and Facebook–postings. All prospective whole blood donors were invited to join the study. A permanent deferral was the only criteria for exclusion from the study. The age limit for blood donation and thus for study participation was 18-70 years (18-59 years for first time donors). Blood samples were taken regardless of the success of the donation. Donation-site nurses actively recruited blood donors to the study only when there was sufficient time for it without impeding the normal donation process. Donors were only recruited in the Helsinki metropolitan area to ensure rapid shipping of samples to the analysis laboratory.

All donations and donor registrations were recorded in a single database (e-Progesa, MAK-SYSTEM). Each participant was asked to fill out a health and lifestyle questionnaire using an internet portal provided at the donation site. The questions are provided in Supplementary Table 1 and are similar to those used e.g. in the Danish blood donor studies [18]. Each participant was asked to fill out the same questionnaire again through the internet portal by an invitation letter sent in June 2018. The internet portal was closed in November 2018. The follow-up questionnaire included 3 additional questions pertaining to quality of life, employment and education. These questions were not asked in the enrollment questionnaire as they were considered too sensitive to be asked at the time of a voluntary and non-remunerated blood donation.

Samples of peripheral blood were collected from the diversion pouch (CompoFlow® Quadruple T&B, Fresenius Kabi, Germany) for successful donations and a separate venous sample was drawn for deferred donations. For blood counts and genomic DNA extraction, samples were collected in 3 ml K2-EDTA tubes (Terumo Europe NV, Belgium) and for CRP, ferritin and sTfR measurements in 3 ml Lithium heparin tubes (Terumo Europe NV, Belgium). A cell pellet was stored for genomic DNA preparation. Samples were sent with the signed informed consent to FRCBS headquarters, located in Kivihaka, Helsinki, Finland, where donor identity and coding on the tubes were double-checked and added to the research database. Plasma samples were sent for laboratory measurements (whole blood count, CRP, ferritin and sTfR; see Supplementary Table 2 for further details) in batches twice a day to the clinical laboratory of the University of Helsinki Central Hospital. Blood counts were measured with Sysmex XE (Sysmex, Japan). The measurements are accredited by the local authority FINAS and the laboratory participates in national and international quality assurance rounds. The tubes were kept in cool storage between transportation and handling and time points from blood draw to analysis were recorded.

### Genotyping

The first 760 enrolled individuals were genotyped using an Illumina HumanCoreExome-24v1-1_A beadchip (Illumina, USA) to estimate ethnic diversity and relatedness. Genotyping of the rest of the participants is under way. See supplementary methods for further detail. The datasets analyzed during the current study are not publicly available due to limitations of the ethical permits which do not allow distribution of personal data, including individual genetic and clinical results.

### Blood count data analysis

The effects of donation time and sampling-to-analysis delay were analyzed using generalized linear multiple regression. Blood count measurement data were used as response variables and donation time, sampling-to-analysis delay, sex and age group (ten year bins) were used as explanatory variables. CRP and ferritin were log2-transformed. Graphical examination of the data suggested interactions between the explanatory factors donation time and sampling-to-analysis delay. Hence, first a model that includes an interaction term for them was tested and if the interaction was not found significant a model without an interaction term was fitted to calculate coefficients of the explanatory variables. The significance of p-values was estimated with Bonferroni correction for multiple testing. The effect of repeated measures was tested by fitting mixed effect models using lme4 R-library [19]. Mixed effect models were identical to the no-interaction models with an additional explanatory variable of individuals as a random effect. Qualitatively similar effect sizes were obtained from the mixed effect models and the no-interaction models. The data were analyzed using R version 3.4.3 [20] and plotted with the ggplot2 R-library [21].

## Results

In total 2584 individual donors (1015 men, 1569 women) gave consent to participate in the study and donated at least one sample. This represent 6% of all donors who visited the study sites during the study. In total, participants donated 8003 samples. None of the participants has withdrawn his or her consent so far (07/2019). 159 participants were new donors (Table 1).

**Table 1.** Characteristics of the study population. Comparison to national (All) and study recruitment site (Study sites) donor populations during the timespan of the study. To facilitate comparison to other cohorts, the count of donations has been normalized from the study period (2.5 years) to 2 years.

|  | Men |  |  | Women |  |  |
| --- | --- | --- | --- | --- | --- | --- |
|  | Study | Study sites | All | Study | Study sites | All |
| n | 1015 | 14850 | 75718 | 1569 | 22735 | 106644 |
| Age (years, median [IQR]) | 49.00 [34.50, 58.00] | 40.00 [28.00, 53.00] | 44.00 [29.00, 56.00] | 43.00 [29.00, 55.00] | 34.00 [25.00, 50.00] | 40.00 [26.00, 53.00] |
| Hb (g/l, mean (sd)) | 152.31 (7.80) | 154.09 (9.33) | 154.86 (10.58) | 138.49 (7.15) | 138.58 (7.96) | 140.15 (9.39) |
| First time donor | 49 ( 4.8) | 3220 (21.7) | 16660 (22.0) | 110 ( 7.0) | 5832 (25.7) | 25378 (23.8) |
| Donation interval (days, median [IQR]) | 107.50 [84.00, 164.25] | 158.50 [85.00, 412.12] | 173.00 [91.00, 461.00] | 162.00 [115.00, 262.25] | 209.00 [107.00, 515.00] | 220.50 [115.00, 564.00] |
| Donations during 2 years (%) |  |  |  |  |  |  |
| 1-3 | 428 (42.2) | 12424 (83.7) | 61830 (81.7) | 1062 (67.8) | 21144 (93.0) | 96910 (90.9) |
| 4-6 | 306 (30.1) | 1542 (10.4) | 8967 (11.8) | 425 (27.1) | 1412 ( 6.2) | 8736 ( 8.2) |
| 7-9 | 215 (21.2) | 754 ( 5.1) | 4145 ( 5.5) | 80 ( 5.1) | 179 ( 0.8) | 998 ( 0.9) |
| 10+ | 66 ( 6.5) | 130 ( 0.9) | 776 ( 1.0) | 0 ( 0.0) | 0 ( 0.0) | 0 ( 0.0) |
| Blood type (%) |  |  |  |  |  |  |
|  | 0 ( 0.0) | 2 ( 0.0) | 14 ( 0.0) | 0 ( 0.0) | 1 ( 0.0) | 15 ( 0.0) |
| A- | 69 ( 6.8) | 897 ( 6.0) | 4468 ( 5.9) | 121 ( 7.7) | 1503 ( 6.6) | 6625 ( 6.2) |
| A+ | 365 (36.0) | 5117 (34.5) | 26852 (35.5) | 546 (34.8) | 7544 (33.2) | 36204 (33.9) |
| AB- | 5 ( 0.5) | 128 ( 0.9) | 854 ( 1.1) | 19 ( 1.2) | 253 ( 1.1) | 1233 ( 1.2) |
| AB+ | 70 ( 6.9) | 881 ( 5.9) | 4549 ( 6.0) | 64 ( 4.1) | 1247 ( 5.5) | 6419 ( 6.0) |
| B- | 21 ( 2.1) | 400 ( 2.7) | 1905 ( 2.5) | 49 ( 3.1) | 566 ( 2.5) | 2799 ( 2.6) |
| B+ | 134 (13.2) | 1972 (13.3) | 10555 (13.9) | 191 (12.2) | 3089 (13.6) | 14836 (13.9) |
| O- | 72 ( 7.1) | 1134 ( 7.6) | 4856 ( 6.4) | 132 ( 8.4) | 1767 ( 7.8) | 7365 ( 6.9) |
| O+ | 279 (27.5) | 4319 (29.1) | 21665 (28.6) | 445 (28.4) | 6765 (29.8) | 31148 (29.2) |
| BMI (cm<sup>2</sup>, mean (sd)) | 25.83 [23.91, 28.31] |  |  | 24.62 [22.31, 28.12] |  |  |
| Weight (kg, mean (sd)) | 85.66 (14.60) |  |  | 71.77 (14.70) |  |  |
| Height (cm, mean (sd)) | 179.82 (6.26) |  |  | 166.64 (5.92) |  |  |

Summary statistics of the age, haemoglobin (Hb) levels, donation interval, count of donations and blood types in the study population were compared to those of the entire national donor population (Group “All” in Table 1 and Supplementary Figures 1 and Figure 2) and to those of the donor population of the recruitment sites (Group “Study Sites” in Table 1 and Supplementary Figures 1 and 2). These three groups were found to differ. The median age of study participants in both sexes was higher than both the national and recruitment site populations, but closer to the national population. The age distribution was bimodal and again the study population resembled more the national donor population than the donor population of the study sites (Supplementary Figure 1A).

### The median donation interval during the study for both sexes was lower than in the national and recruitment site populations

Accordingly, donation counts were higher (Table 1 and Supplementary Figure 2). Mean Hb (g/l) was lower in men in the study population than in the national and recruitment site populations (Table 1 and Supplementary Figure 1B). However, in women, mean Hb (g/l) in the study population was higher than in the recruitment site population, which reflects the higher age of the study population. Consequently, the study population consisted of a markedly higher proportion of older donors with frequent donation activity and fewer new donors. The distribution of blood types between different populations resembled each other closely except that the study population has been more comprehensively blood typed due to their frequent donor status (Table 1).

### The during-study donation attempt frequency was found to be higher than the before-study donation attempt frequency for study participants

(median of during-study – before-study frequency = 0.6, p < 2e-44 for men; 0.6, p< 9e-80 for men, Wilcoxon signed rank test, Figure 1). For each donor, the before-study donation frequency was calculated as the annual average count of donations for the two years before study enrollment. The during-study frequency was calculated as the annual average count of donations from the time of enrollment to the study end. Donors who enrolled after 2016.12.08 (less than one year before the end of the study) and new donors were excluded. 2271 (906 men and 1365 women) donors were therefore included in the analysis.

**Figure 1:**
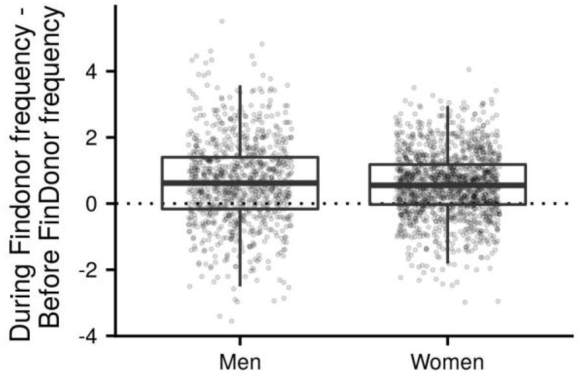
Difference of during study donation attempt frequency and before study donation attempt frequency. Before study donation frequency for a donor was calculated as the annual average count of donations from two years before the study enrollment.

Of the 2584 study participants, 2562 (99%; 1004 men, 1558 women) answered the enrollment questionnaire and 1477 (57%; 597 men, 880 women) the follow-up questionnaire. We used self-reported health as the main indicator of blood donor health. Donors answered the question “How would you rate your recent health in general” on a five-point scale (Figure 2). At the population-level, the distributions of answers given in the enrollment questionnaire (filled at time of donation) and in the follow-up questionnaire (answered outside donation visits) are similar. At the individual level, there did not seem to be any systematic increase or decrease of participants’ health ratings during the study period.

**Figure 2:**
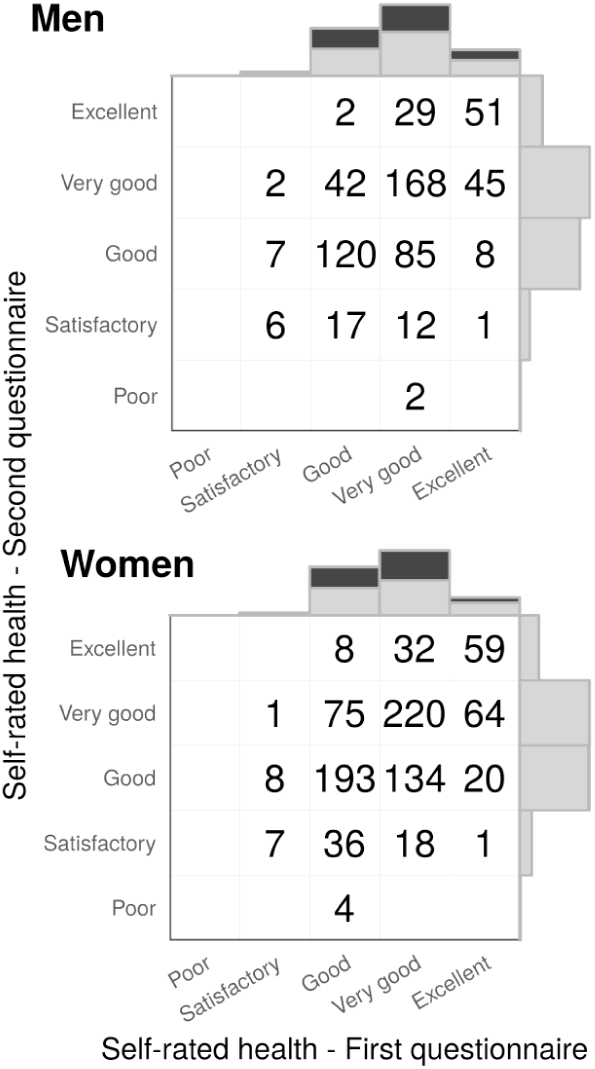
Self-rated health in study enrollment and follow-up questionnaire. Marginal distributions show the total amount of answers and numbers in cells the specific combinations of answers to the two questionnaires.

We asked donors about their level of education and their employment situation to assess any potential effects of these factors on self-reported health (Figure 3). Most of the donors who replied to both questionnaires had at least a vocational college education (men: 59 %, women: 57 %). Furthermore, a majority of these donors were either employed full-time (men: 70 %, women: 67 %) or pensioners (men: 17 %, women: 14 %).

**Figure 3:**
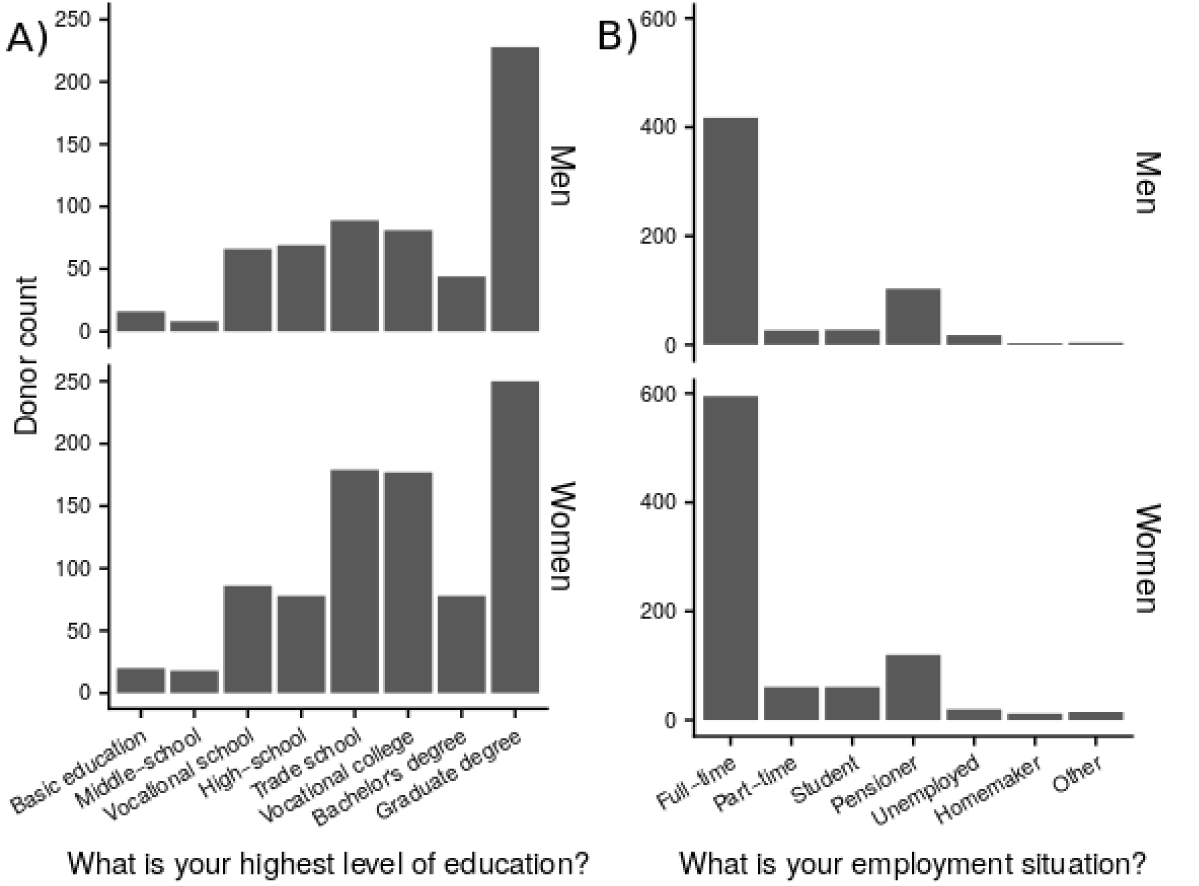
Distributions of education (A) and employment (B) status among participants who answered the follow-up questionnaire.

At least two blood samples could be collected from 1976 donors (N= 840 men, N= 1136 women, see Supplementary Table 3 for further details).

DNA samples from 760 participants were subjected to a genome-wide SNP array screen. To demonstrate the genetic homogeneity of the study population, we performed principal component analysis (Supplementary Figure 3) that demonstrated that all 760 participants could be classified as Northern Europeans.

For each blood-count test, the sampling time (Figure 4) as well as the time when the analysis result was ready were entered to the laboratory information system. The difference between these two time points was used as the sampling-to-analysis delay time (Figure 5). To assess whether the trends shown in Figures 4 and 5 were significant, a multiple regression model was fitted for each individual blood count measurement including both donation time, sampling-to-analysis delay, an interaction term between these two variables, and sex and age as explanatory factors (Table 2). The interaction term was used to evaluate if the two time variables could be separated from each other in this cohort. If the interaction term was not found to be significant then the coefficient and p-value of the time variables were calculated from a model without the interaction term. While the coefficient from a model without the interaction term is the unit change in an hour, e.g. Hb drops on average 0.2 g/l per hour when going from 8 a.m. to 8 p.m., there is no such intuitive interpretation for coefficients from a model with an interaction term. An effect in same direction and very similar size range has been shown previously from finger prick Hb measurements [22].

**Table 2.** Significance of explanatory variables for each measurement. For each variable a model was fitted with age, sex, sampling-to-analysis delay (h) and donation time (h) as explanatory variables. A model including an interaction term was first fitted (“p-value from a model with interaction”). If the interaction term was not significant, coefficient and p-value is provided for a model without interaction. P-values that are significant after Bonferroni correction of alpha (original alpha 0.05) are marked with an asterix (*).

| Measurement | Number of measurements | Mean | Coefficient from a model without interaction |  | p-value from a model without interaction |  | p-value from a model with interaction |  |  |
| --- | --- | --- | --- | --- | --- | --- | --- | --- | --- |
|  |  |  | Sampling-to-analysis delay (h) | Donation time (h) | Sampling-to-analysis delay (h) | Donation time (h) | Sampling-to-analysis delay (h) | Donation time (h) | Sampling-to-analysis delay : Donation time |
| log2(CRP mg/l) - C-Reactive Protein | 7906 | 3.33 | 1.000 | 0.999 | 7.2E-01 | 6.2E-01 | 3.1E-01 | 2.8E-01 | 3.2E-01 |
| Eryt (x 10<sup>12/l</sup>) - Erythrocyte count | 7909 | 4.79 | -0.001 | -0.011 | 7.9E-03 | 9.7E-11 * | 1.4E-01 | 8.9E-01 | 6.8E-02 |
| log2(Ferritin µg/l) - Ferritin | 7902 | 37.63 |  |  |  |  | 2.6E-04 * | 2.1E-04 * | 1.1E-03 * |
| Hb (g/l) - Haemoglobin | 7910 | 142.71 | -0.031 | -0.178 | 2.7E-02 | 7.6E-05 * | 2.9E-01 | 7.8E-01 | 1.9E-01 |
| HKR (%) - Hematocrit | 7919 | 42.72 | 0.028 | -0.083 | 2.1E-14 * | 8.7E-12 * | 2.2E-01 | 1.3E-04 * | 3.0E-02 |
| Leuk (x 10<sup>9/l</sup>) - Leukocyte count | 7909 | 6.60 | 0.004 | 0.108 | 9.3E-02 | 2.6E-41 * | 3.1E-02 | 1.1E-07 * | 4.9E-02 |
| MCH (pg / cell) - Mean corpuscular hemoglobin | 7918 | 29.92 | 0.003 | 0.028 | 2.6E-01 | 1.4E-03 * | 3.4E-01 | 8.0E-01 | 2.7E-01 |
| MCHC (g/l) - Mean corpuscular hemoglobin concentration | 7910 | 334.31 |  |  |  |  | 8.4E-06 * | 6.9E-15 * | 2.8E-12 * |
| MCV (fl) - Mean corpuscular volume | 7914 | 89.38 |  |  |  |  | 3.0E-04 * | 4.1E-06 * | 2.0E-07 * |
| RDW % - Red cell Distribution Width | 7920 | 13.88 |  |  |  |  | 7.8E-03 | 1.3E-04 * | 1.5E-03 * |
| sTfR (mg/l) - soluble Transferrin Receptor | 7646 | 3.93 |  |  |  |  | 2.1E-04 * | 5.8E-05 * | 4.8E-05 * |
| Trom (x 10<sup>9/l</sup>) - Thrombocyte count | 7911 | 247.21 | 0.240 | 0.430 | 1.5E-03 * | 8.0E-02 | 9.7E-01 | 9.8E-01 | 6.7E-01 |

**Figure 4:**
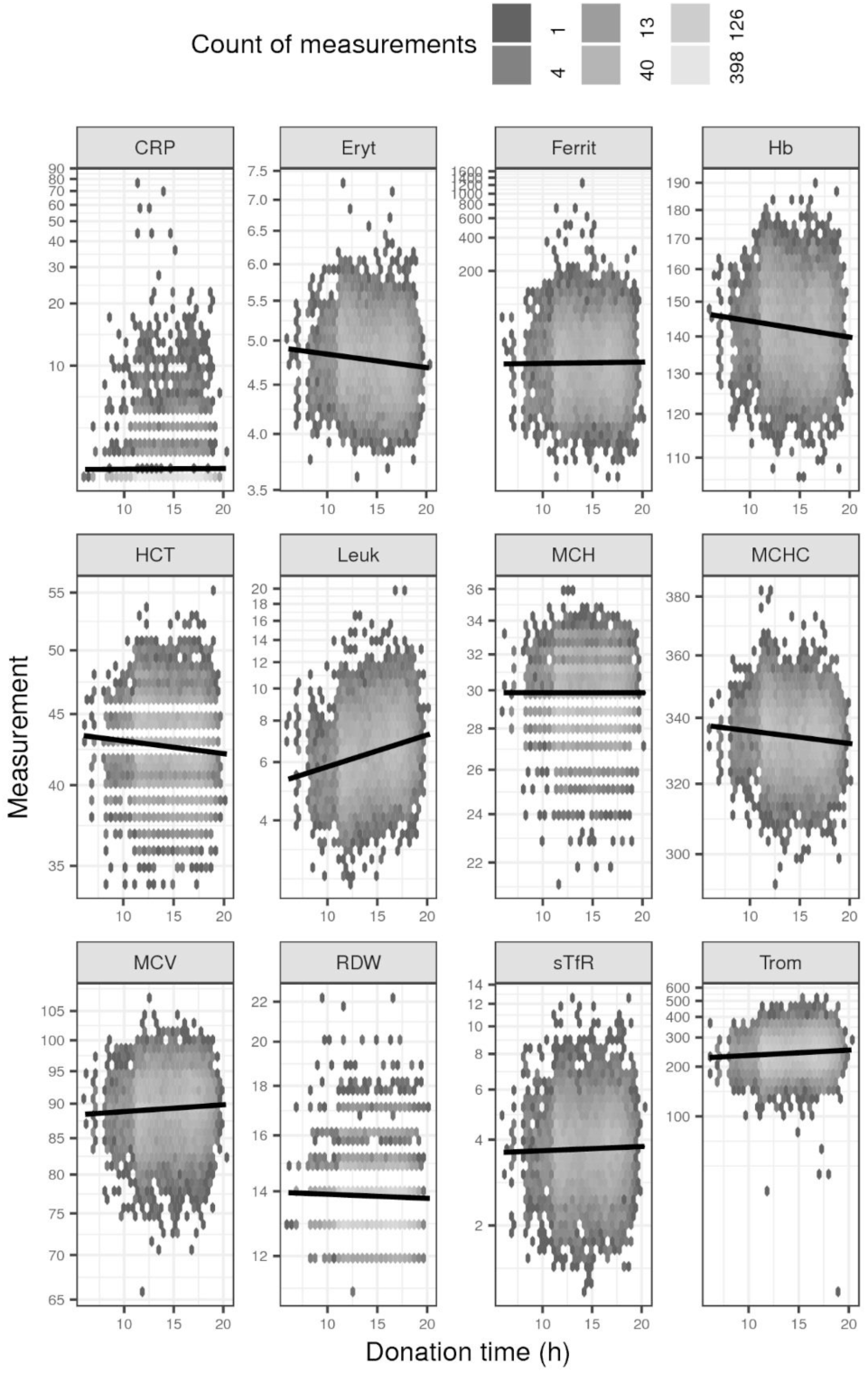
Effect of donation time to measurement values. Data is binned to hexagons and the color of each hexagon shows how many individual measurements are contained in each hexagon. The black line shows a linear regression trend line. See Table 2 for full description of measured variables.

**Figure 5:**
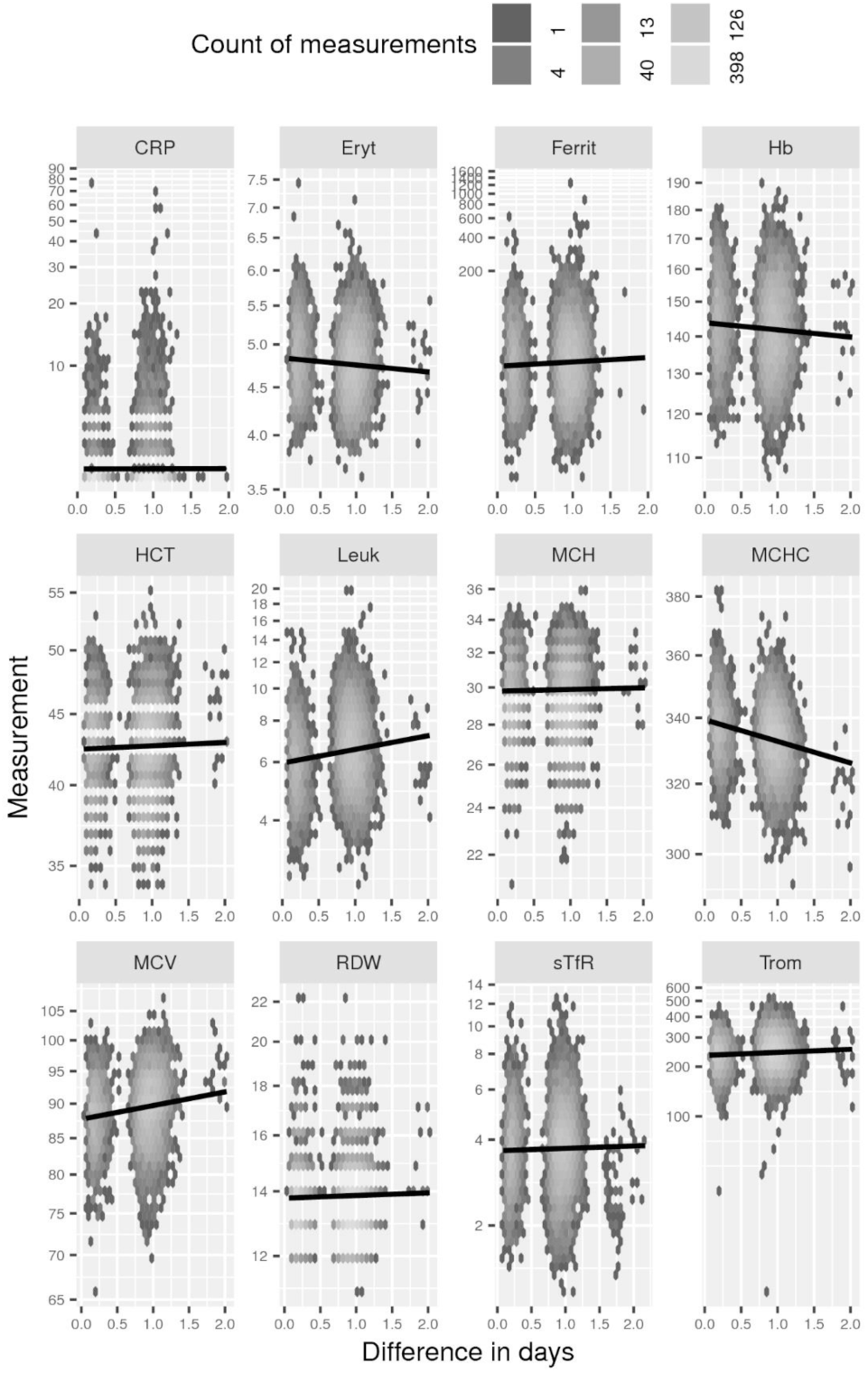
Effect of sampling-to-analysis delay time (d) to measurement values. See Figure 4.

### For hematocrit and platelet count there was a separable significant sampling-to-analysis delay time (h) effect and for erythrocyte count, Hb, hematocrit, MCH and leukocyte count there was a separable significant donation time (h) effect

For ferritin, MCV, MCHC, RDW and sTfR there were significant interactions of donation time (h) and sampling-to-analysis delay time (h) and hence their individual effects could not be separated.

A single elevated CRP measurement, i.e. a value above 10 that may point to inflammation, was found in 1.8 % of women and 0.3 % of men. Respectively, 14.3 % and 7.6 % had a single measurement between 3 and 10, i.e. a low grade inflammation. To investigate if these measurements could point to chronic inflammation we investigated the data graphically (Supplementary Figure 4). No donor was found to have two CRP measurements above 10, hence, they most likely had no chronic inflammation.

## Discussion

The present cohort was collected primarily for studies investigating the effects of regular blood donation on donors’ health, in particular regarding iron stores and factors regulating them in a population with a long-standing systematic iron replacement policy. The cohort complements those collected elsewhere [12–14,18]. We put special emphasis on careful collection and follow-up of plasma and serum samples to understand possible confounding factors in measurements of iron markers. Modelling blood count and related data is not simple as HCT, MCHC, MCV, RDW and CRP show properties of counts instead of true continuous measurements, hence, defying the assumptions of standard linear regression. To mitigate this, we experimented with various transformations, link functions and error distributions in the modelling process to no avail. The results (Figures 4 and 5, Table 2) clearly demonstrate that it is essential to know the detailed conditions of collected samples if accurate results are to be assumed. Recent literature suggests that the stability of blood count parameters in cool storage is process- and device-specific, hence it should be addressed in every cohort specifically [23,24]. There are interesting studies demonstrating large effects related to the time of drawing the samples [25] and this variation also is relevant in everyday blood donation practice [22]. However, for many screening purposes the variation in measurements found in the present study is apparently not critical.

Another factor that is essential to know in cohort studies is how well the study population represents the overall target population, here blood donors. Based on the demographics, Hb distribution and donation activity, participants (Supplementary Figures 1 and 2, and Table 1) of the FinDonor 10 000 cohort can be regarded in general to represent Finnish donor pool despite some bias toward frequent and committed donors. As a key question to be clarified with the cohort is related to effects of frequent donation, this bias may not have serious drawbacks for our future studies.

In Finland all blood donations are voluntary and non-remunerated. Such a blood donation system could be sensitive to any additional burden to donors (e.g., participating to research). However, as suggested by two previous studies [13,26], it was found that the donation frequency of study participants did not decrease during the study in comparison to their previous donation activity (Figure 1). It is of note that no additional efforts to remind participants to donate were made. Hence, carrying out blood donation research of this scale embedded in the regular blood donation process of FRCBS is clearly feasible.

Self-reported health is one of the most widely used instruments to assess general health in public health studies and associates strongly and constantly with mortality [27]. When a person arrives to donate she considers herself healthy. Hence, answers to questions regarding her health status at donation time could be biased towards feeling healthier than she generally feels. To check for the presence of this possibly important source of bias, we asked donors to answer the study questionnaire again at the end of the study, outside any blood donation visit. Based on the comparison of the enrollment and follow-up questionnaires, it seems that this bias, if present at all, is negligible in our cohort.

The socio-economic make-up of blood donor populations vary amongst countries and cultural contexts [28]. As socio-economic background is also an important determinant of health, it was important to assess the educational background and employment situation of our specific donor population. In the FinDonor cohort’s follow-up questionnaire, university graduates (38% of men and 28% of women) and full-time employees (70% of men and 67% of women) are overrepresented in comparison to the general Finnish population. In the general Finnish population of at least 15 years of age in 2017, only 10 % had at least a graduate level degree and only 38 % were employed full-time [https://findikaattori.fi/]. Our data may however be affected by a self-selection bias. Answering the follow-up questionnaire required the donor to access the internet portal from their own device, therefore more educated donors and/or tech-savvy donors might have been more likely to answer it.

The prevalence of CRP between 3 and 10 was very similar to what has been reported in the Danish blood donor population (14.4% women and 6.1% men [29]). The prevalence of CRP > 10 was found to be below 2%. Approximately 7% of the apparently healthy Finnish adults have earlier been reported to have CRP levels over 10 [30,31]. As a minor CRP elevation (between 3 and 10) does not directly point to inflammation but rather to various mild tissue stress or injury [32], the FinDonor 10 000 cohort appears to have a markedly low level of chronic inflammation. This is in accordance with the healthy donor effect, [33] i.e. healthier individuals are selected as donors and are able to maintain the habit from years to decades.

## Conclusions

Together with similar cohorts from other populations [12–14,18] the FinDonor cohort provides us with tools to identify the potential donor groups at risk for iron deficiency and the genetic and non-genetic factors associated with this increased risk, as well as to study other health effects of blood donation. These background facts are needed for ensuring safer and more personalized blood donation in the future.

## Acknowledgements

We want to thank the nurses and other staff of the Kivihaka, Sanomatalo and Espoo donation sites for donor recruitment, blood donors for participation in the study and Veera Raivola for help with sociological issues.

## Author contributions

Study concept: JP and JC;

Management and planning of recruitment and sample processing: PM, PN, EP and NN;

Data processing and analysis: AL, ML and MA;

Interpretation of results: PN, ML, EP, JC, MA, JP;

Manuscript preparation: JP and MA.

