## Supplemental tables, figures and methods for "FinDonor 10 000 study: A cohort to identify iron depletion and factors affecting it in Finnish blood donors"

Short title: Finnish blood donor cohort


Lobier, Muriel

Niittymäki, Pia

Nikiforow, Nina

Palokangas, Elina

Larjo, Antti

Mattila, Pirkko

Castrén, Johanna

Partanen, Jukka*

Arvas, Mikko*

Finnish Red Cross Blood Service, Research and Development. Kivihaantie 7, 00310 Helsinki, Finland.


**Supplementary Table 1** is provided as a separate file.

| | 18th May 2015 to 31st May 2016 | | | 1st June 2016 onwards | | |
|---|---|---|---|---|---|---|
| | reagent | device | reagent manufacturer | reagent | device | reagent manufacturer |
| CRP | CRPL3 | Roche Modular | Roche Diagnostics | CRP VARIO | Abbott Architect | Abbott |
| Ferritin | FERR4 | Roche Modular | Roche Diagnostics | Abbott Architect Ferritin | Abbott Architect | Abbott |
| sTFR | Tina-quant Soluble Transferrin Receptor | Roche Modular | Roche Diagnostics | Tina-quant Soluble Transferrin Receptor | Abbott Architect | Roche Diagnostics |

**Supplementary Table 2**. Reagents and devices used for CRP, ferritin and sTfR measurements.

| Gender | Count of study donation attempts | Count |
|---|---|---|
| Women | 1 | 433 |
| Women | 2 | 407 |
| Women | 3 | 316 |
| Women | 4 | 196 |
| Women | 5 | 124 |
| Women | 6 | 65 |
| Women | 7 | 22 |
| Women | 8 | 6 |
| Men | 1 | 175 |
| Men | 2 | 194 |
| Men | 3 | 175 |
| Men | 4 | 134 |
| Men | 5 | 121 |
| Men | 6 | 80 |
| Men | 7 | 60 |
| Men | 8 | 35 |
| Men | 9 | 20 |
| Men | 10 | 12 |
| Men | 11 | 5 |
| Men | 12 | 4 |
| | Count of donors | 2584 |

**Supplementary Table 3** Counts of study donation attempts during which study blood sample was given.

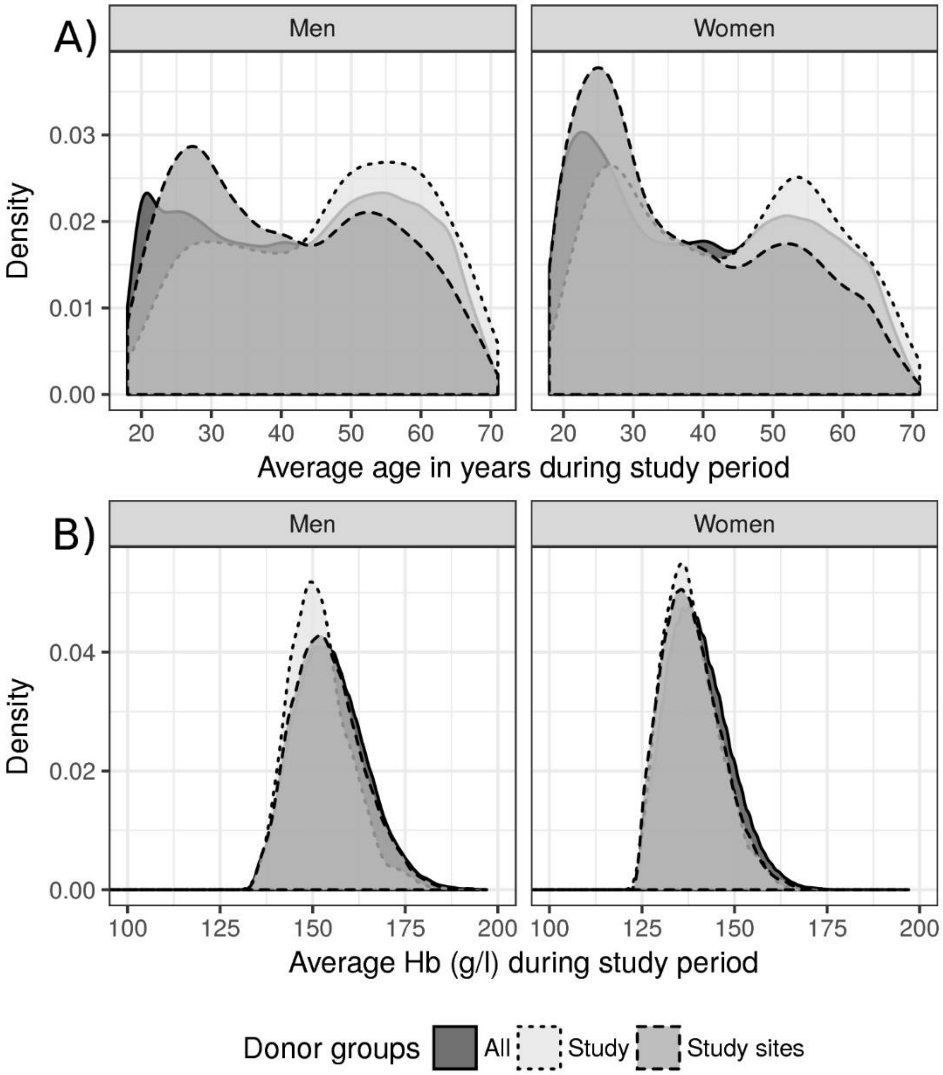


**Supplementary Figure 1: Age and Hb distributions of study participants.** Density function of A) age distributions of all donors ("All"), all donors that donated in the collection sites where study took place ("Study sites") and study participants ("Study") during the study period. Respectively the density distribution of B) hemoglobin. Average over all donation events during the study period is used as the value of age and hemoglobin. Mean of hemoglobin was "All": 140 women and 154 men; "Study sites" 138 and 154 and "Study": 138 and 151.

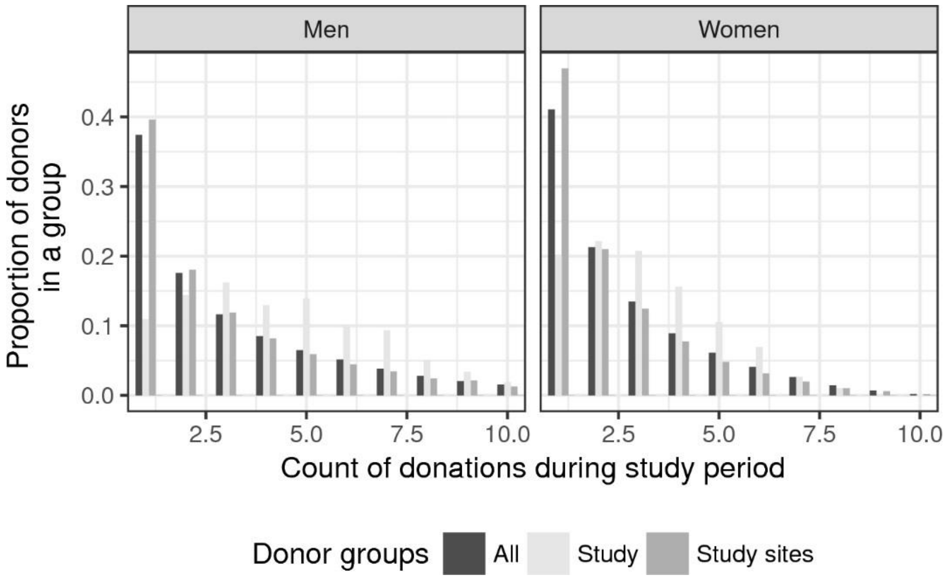


**Supplementary Figure 2**: **Counts of donations of study participants during study period.** "All" are donations by all donors that donated during the study at any FRCBS site. "Study sites" are donations of all donors in the collection sites where study took place. "Study" are study participants. Y-axis shows the proportion of the donors in the group (All, Study sites, Study) that donated a certain number of donations during the study period (on x-axis).

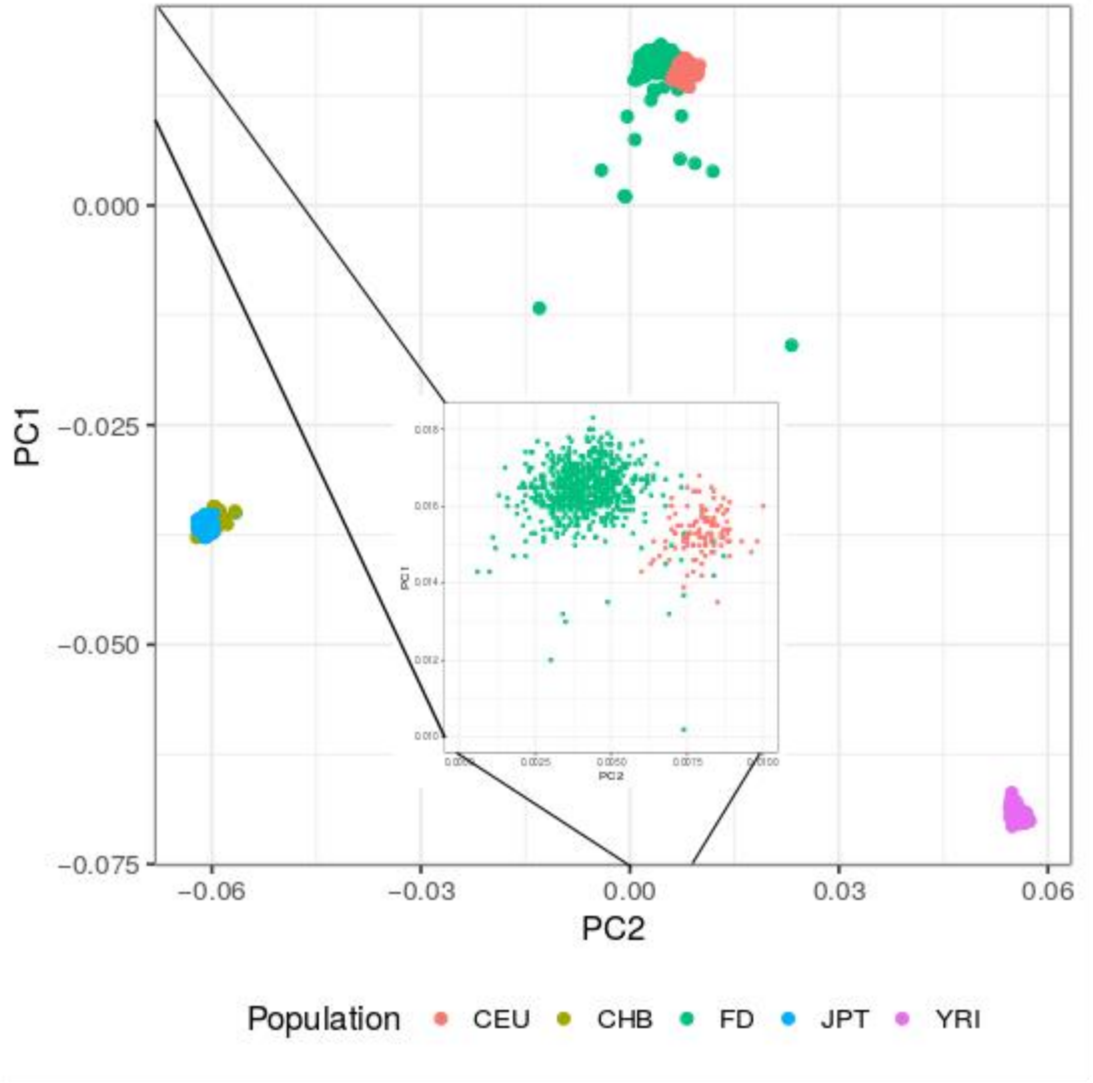


**Supplementary Figure 3: Relatedness of 760 FinnDonor participants and selected 1000 Genome reference populations.** Principal component analysis of similarity of exome genotyping results. Populations: CEU = Utah Residents (CEPH) with Northern and Western European Ancestry, CHB = Han Chinese in Beijing, China, FD = FinnDonor, JPT = Japanese in Tokyo, Japan, YRI = Yoruba in Ibadan, Nigeria. Inset shows the separation of FinnDonors from the CEU.

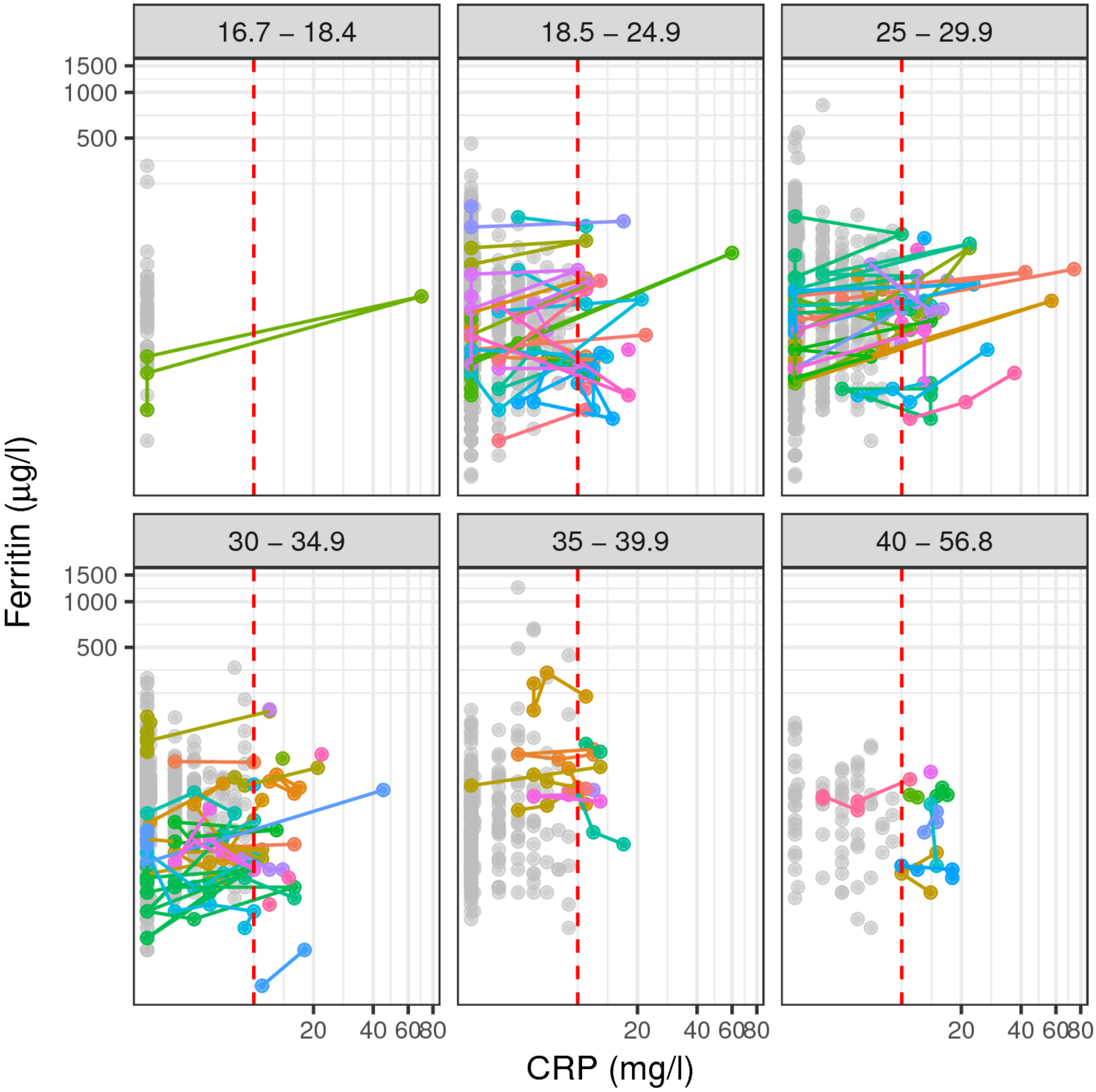


**Supplementary Figure 4: Ferritin versus CRP divided by BMI groups.** Each dot is a single blood sample. For individuals which have at least one sample with CRP > 10 mg/l (red dotted vertical line) dots are colored in the same color and connected by a line. Due to lack of distinguishable colors, colors are recycled from an individual to another. To aid visual separation of individuals the data is divided into 6 plots by WHO BMI based obesity classes.

### Supplementary methods

Genotyping was performed at the FIMM Technology Centre, Helsinki, Finland. DNA samples from the Finnish cohort were extracted using the QIAamp DNA Blood Mini Kit (Qiagen, Germany). Imputation was carried out as in [1] according to instructions of [2] using plink 1.90b3.29 [3] for quality filtering and IMPUTE2 [4] for the actual imputation with 1000 Genomes Phase 3 [5] as a phased reference. Post-imputation filtering excluded variants having an IMPUTE2 INFO-field <0.5. After post-imputation filtering 11 880 443 variants were included. PCA (Principal Component Analysis) of the imputed and filtered dataset was carried out with eigensoft version 6.1.4 [6].
